# A PCR amplicon–based SARS-CoV-2 replicon for antiviral screening

**DOI:** 10.1101/2020.08.28.267567

**Authors:** Tomohiro Kotaki, Xuping Xie, Pei-Yong Shi, Masanori Kameoka

## Abstract

The development of specific antiviral compounds to SARS-CoV-2 is an urgent task. One of the obstacles for the antiviral development is the requirement of biocontainment because infectious SARS-CoV-2 must be handled in a biosafety level-3 laboratory. Replicon, a non-infectious self-replicative viral RNA, could be a safe and effective tool for antiviral screening; however, SARS-CoV-2 replicon has not been reported yet. Herein, we generated a PCR-based SARS-CoV-2 replicon. Eight fragments covering the entire SARS-CoV-2 genome except S, E, and M genes were amplified with HiBiT-tag sequence by PCR. The amplicons were ligated and *in vitro* transcribed to RNA. The cells electroporated with the replicon RNA showed more than 3,000 times higher luminescence than MOCK control cells at 24 hours post-electroporation, indicating robust viral translation and RNA replication. The replication was drastically inhibited by remdesivir, an RNA polymerase inhibitor for SARS-CoV-2. The IC_50_ of remdesivir in this study was 0.29 μM, generally consistent to the IC_50_ obtained using infectious SARS-CoV-2 in a previous study (0.77 μM). Taken together, this system could be applied to the safe and effective antiviral screening without using infectious SARS-CoV-2. Because this is a transient replicon, further improvement including the establishment of stable cell line must be achieved.

## Introduction

Severe acute respiratory syndrome-coronavirus 2 (SARS-CoV-2) has been causing a catastrophic pandemic worldwide. The symptoms of SARS-CoV-2 infection (coronavirus disease 2019 [COVID-19]) ranges from asymptomatic to fever, acute respiratory distress, pneumonia, and ultimately death [1]. To date, several antiviral drugs such as remdesivir (viral RNA–dependent RNA polymerase [RdRp] inhibitor for Ebola virus) have been repurposed for COVID-19 therapy [2]. Nevertheless, the mortality was still high (above 5%) [3]. Therefore, it is important to develop antiviral agents that can specifically inhibit the propagation of SARS-CoV-2.

One of the obstacles for the antiviral screening of SARS-CoV-2 is biosafety concern. The high infectivity and mortality of SARS-CoV-2 have rendered antiviral screening difficult. Because SARS-CoV-2 was classified as a biosafety level-3 (BSL-3) pathogen, it must be handled in a BSL-3 laboratory. The construction of a safe and high throughput antiviral screening system has been coveted.

The replicon system could be a useful tool for safe and efficient antiviral screening. Replicon is a non-infectious, self-replicative RNA that lacks the viral structural genes and retains the genes necessary for RNA replication [4,5]. Because the replicon lacks viral structural genes, infectious virions are not produced from the transfected cell, thus reducing the biosafety concern. Additionally, the insertion of reporter gene into the replicon genome enables us to easily monitor the viral replication. The construction of a replicon system would accelerate the antiviral development.

SARS-CoV-2 belongs to the genus *betaoronavirus* of the family *coronaviridae* [6]. The genome of coronaviruses is single-stranded RNA ranging from 27 to 32 kb, the largest of any other known RNA viruses. Its large genome size and the existence of bacteriotoxic elements hindered the generation of reverse genetic systems and replicon. Several strategies have been adopted to overcome this obstacle: multiple plasmid system followed by *in vitro* DNA ligation or single bacterial artificial chromosome (BAC) plasmid system [7–9]. With these strategies, the infectious clones of SARS-CoV-2 and its reporter variants have been developed [10]. However, for now, SARS-CoV-2 replicon has not been reported elsewhere.

Herein, we generated a first SARS-CoV-2 replicon by the *in vitro* ligation of PCR amplicons. The results demonstrated its use for antiviral screening without using the infectious SARS-CoV-2 virion.

## Results

### The construction of a SARS-CoV-2 replicon

We took an *in vitro* ligation strategy, similar to that used for constructing a SARS-CoV-2 infectious clone [10] (Figure 1A, B). The genome of replicon included viral non-structural proteins (encoded in open reading frame [ORF]1a and 1b) and N protein that were required for RNA replication. Meanwhile, the viral structural proteins (S, E, and M) were excluded so as not to produce infectious virion. For facilitating the detection of viral protein, HiBiT-tag was incorporated into the C-terminus of N protein. SARS-CoV-2 5’ untranslated region (UTR), ORF1a, and 1b were separately amplified in the fragment 1 (F1) to F7. Then, N (including the closest transcription regulatory sequence [TRS] on 5’ upstream: ACGAACAAACTAAA), HiBiT-tag, and 3’UTR were amplified in the F8. Each amplicon comprised the BsaI recognition sites at the both 5’ and 3’ termini. Figure 1C shows the detailed information of the fragments.

**Figure 1.**
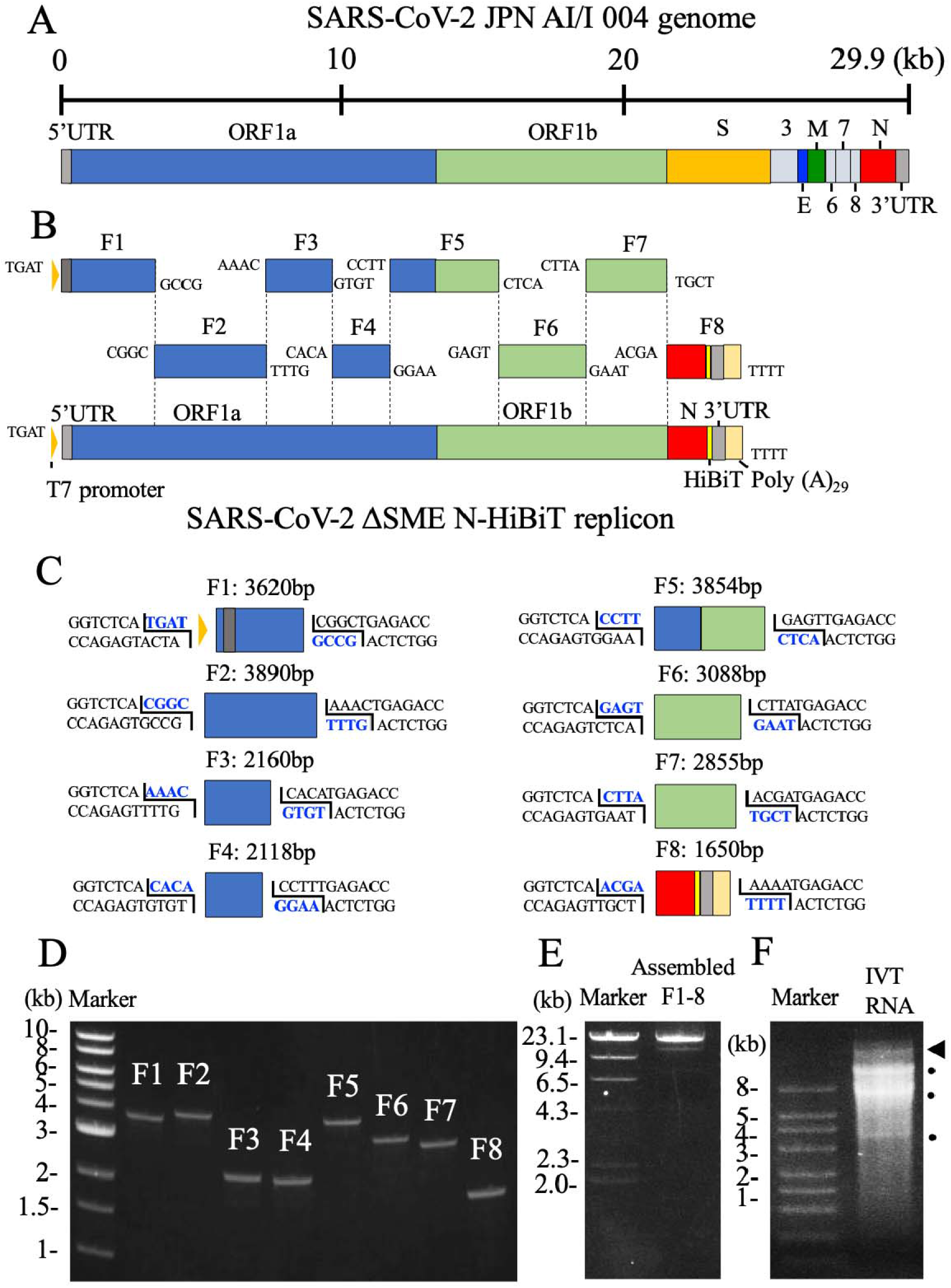
Construction of a SARS-CoV-2 replicon. (A) Genome structure of SARS-CoV-2. The untranslated regions (UTRs), open reading frames (ORFs), and structural proteins (S, E, M, and N) are indicated in this figure. (B) Strategy for the *in vitro* assembly of a SARS-CoV-2 replicon DNA. The nucleotide sequences of the overhang are indicated in this figure. The replicon DNA was assembled using *in vitro* ligation. (C) Detailed terminal sequences of each DNA fragment. Both 5’ and 3’ terminal sequences were recognized by BsaI. The overhang sequences were shown in blue. (D) Electrophoresis of the eight DNA fragments. Eight purified DNA fragments (about 100 ng) were run on a 1.0% agarose gel. The 1-kb DNA ladders are indicated in this figure. (E) Electrophoresis of an assembled DNA. About 200 ng of assembled DNA was run on a 1% agarose gel. The λ-HindIII digest marker is indicated in this figure. Successfully assembled replicon DNA was 23.2kb. (F) Electrophoresis of RNA transcripts. About 1 μg of *in vitro* transcribed (IVT) RNAs were run under denaturing conditions. RNA ladders are indicated in this figure. The triangle indicates the genome-length RNA transcript (23kb), whereas the circles show the shorter RNA transcripts. Because the biggest size of RNA marker was 8 kb, the estimation of the size of RNA transcripts was not accurate.

The viral RNA extracted from the culture fluid of SARS-CoV-2–infected Vero E6 cell was used as a template for RT-PCR. Table 1 shows the primer sets used for the amplification of above-described eight fragments (Figure 1D). The fragments were assembled in a two-step ligation: (1) all the eight fragments were digested with BsaI, followed by the ligation of two adjacent fragments (e.g. F1 and F2 for F1–2) to produce four assembled fragments; (2) the ligated fragments were gel extracted and mixed, followed by a further ligation to construct the full-length replicon DNA. The size of the successfully ligated replicon DNA was 23.2 kb (Figure 1E). *In vitro* transcription using the replicon DNA produced multiple bands (Figure 1F). Of these bands, the highest band might represent the full-size replicon (indicated by arrow). Because the biggest size of RNA marker was only 8 kb, the estimation of the size of RNA transcripts was not accurate.

**Table 1.**
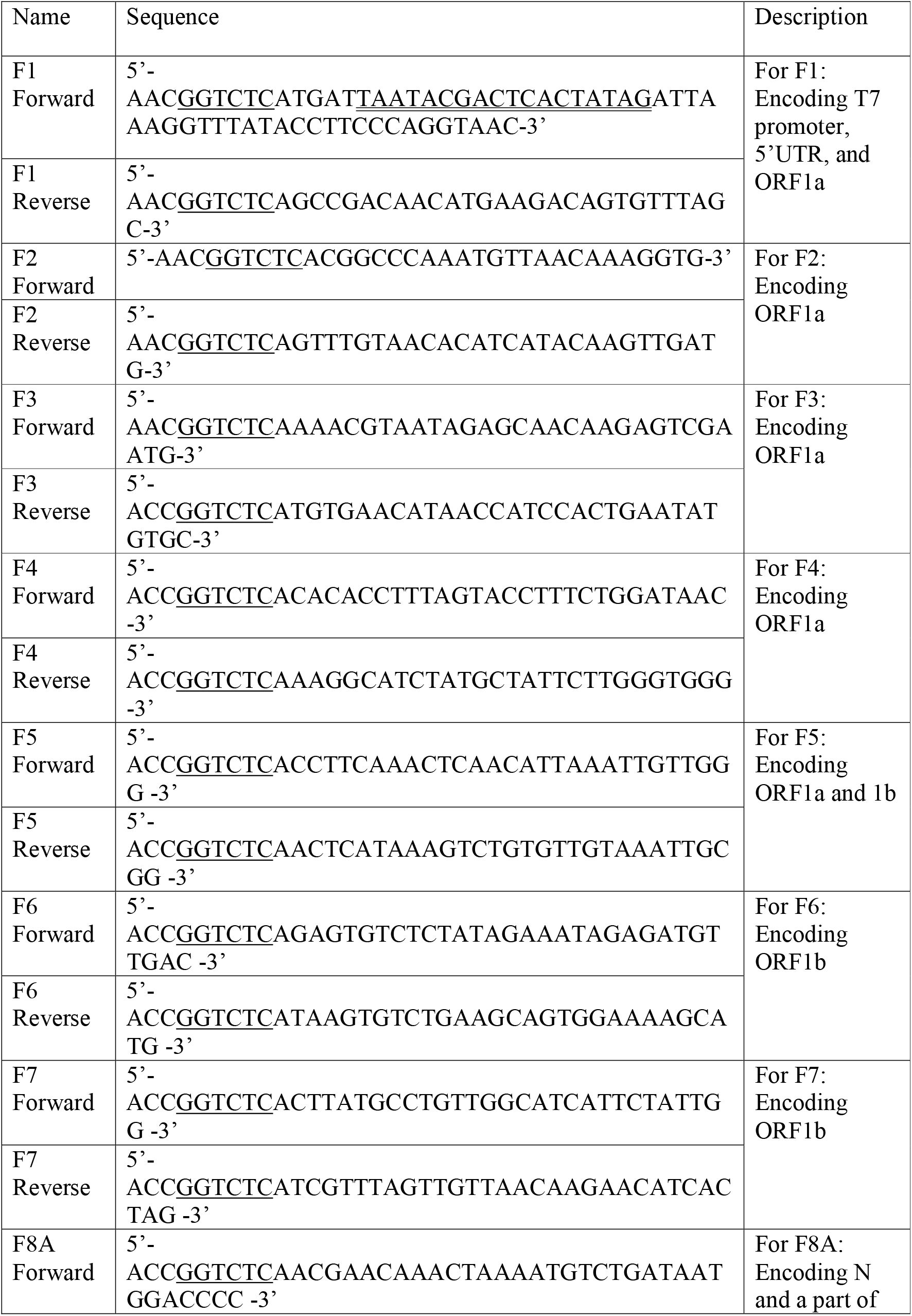
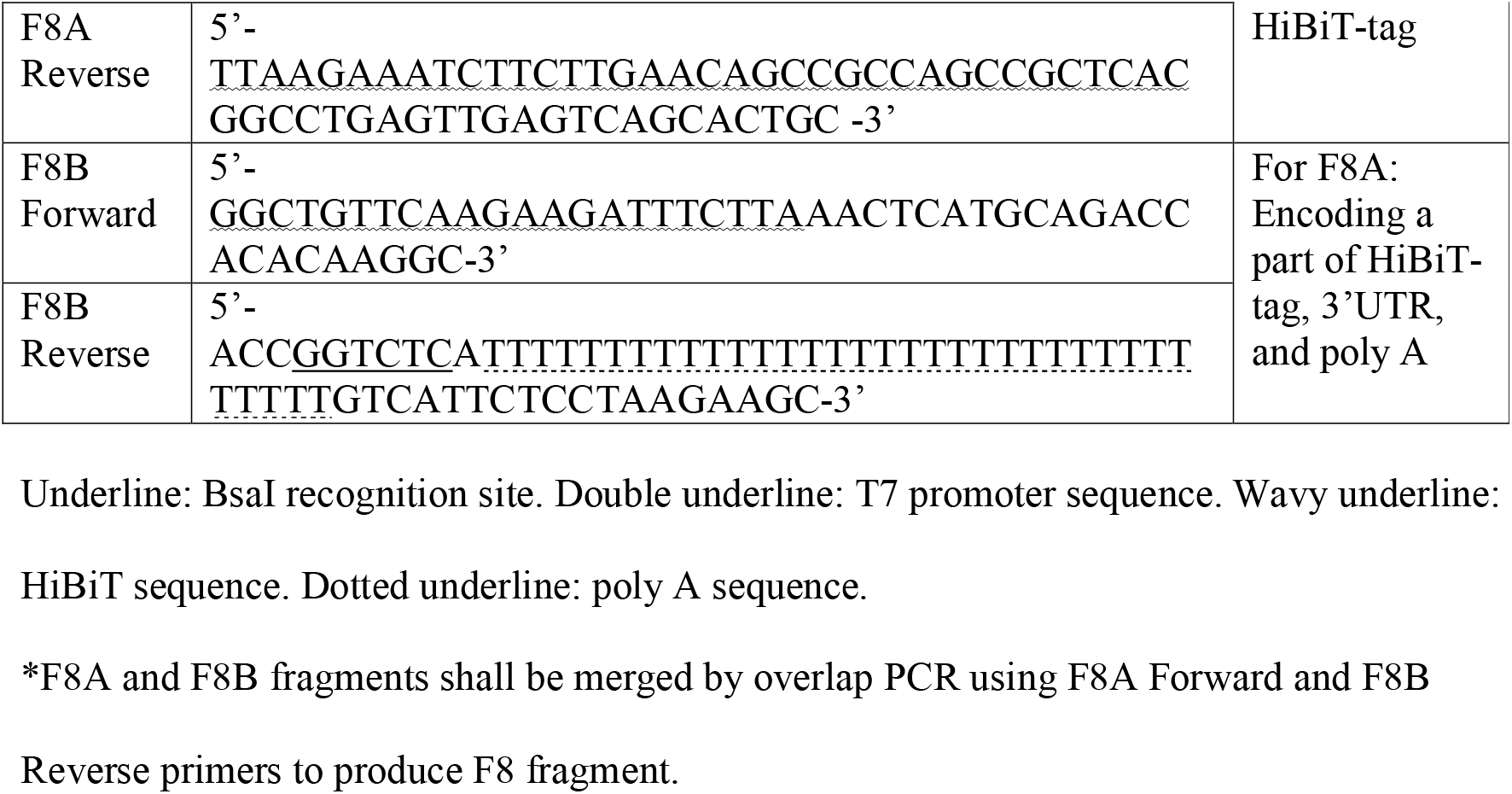
Primer list.

### Characterization of a SARS-CoV-2 replicon

The *in vitro* transcribed RNA was directly electroporated (without gel purification) into BHK-21, HEK-293T, or CHO-K1 cells to determine the most robust replicon system. In BHK-21 and 293T cells, luminescence signals were stable and similar to the MOCK control at two to six hours post-transfection (hpt) (Figure 2A). At 24–48 hpt, the signals increased to 10–100 times. These data implied that the replicon was replicated but was not robust in these cell lines. Meanwhile in the CHO-K1 cell, the signals started to increase as early as 4–6 hpt, indicating replication and the subsequent translation of the replicon (Figure 2A). At 24–48 hpt, the signals increased by more than 3,000 times than the MOCK control. Thus, the CHO-K1 cell was the most suitable cell line for the robust replication of the replicon, and used for the subsequent experiments. The viral N protein and NSP8 (a component of RNA replication complex encoded in ORF1a) expressions were confirmed by immunofluorescent assay (IFA) (Figure 2B, 2C). These data indicated that the replicon was successfully constructed and replicative.

**Figure 2.**
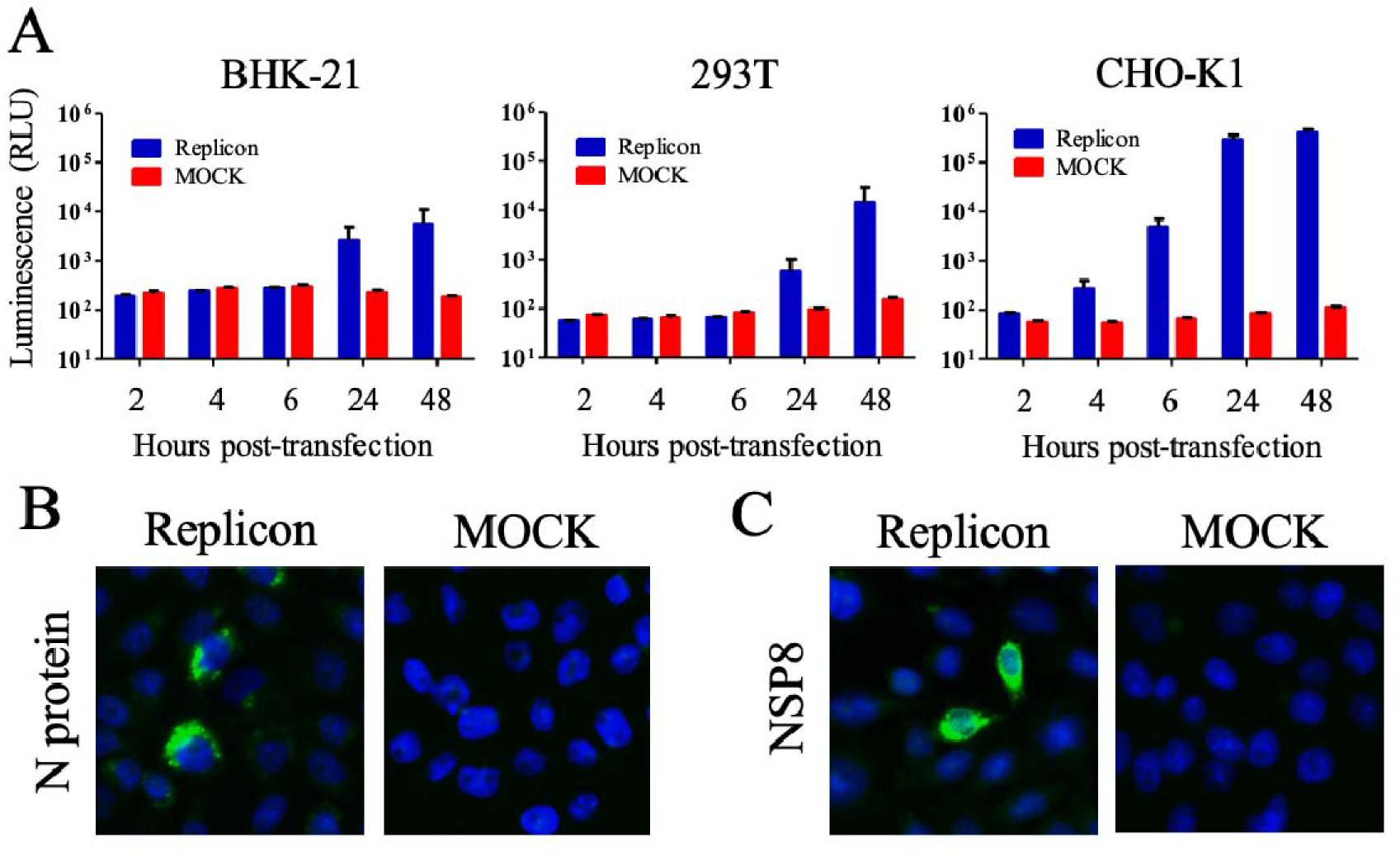
Characterization of a SARS-CoV-2 replicon. (A) Kinetics of luminescence signal. Three cell lines were electroporated with 5 μg of replicon RNA. Intracellular luminescence signals were measured at the indicated time points. The mean and standard error of two independent experiments are shown in this figure. (B) The detection of N protein by IFA. The CHO-K1 cell was electroporated with 5 μg of replicon RNA. The cells were fixed with 4% paraformaldehyde, followed by permeabilization with 0.5% Triton-X. The expression of N protein was detected using anti-N mAb and goat-anti-mouse IgG conjugated with Alexa Fluor 488. Nucleus was stained by DAPI. (C) The detection of NSP8 protein by IFA. The expression of NSP8 protein was detected using anti-NSP8 mAb and goat-anti-mouse IgG conjugated with Alexa Fluor 488.

### Antiviral evaluation

Next, we tested if this RNA replicon could be used for drug screening. Remdesivir, an RdRp inhibitor effective for SARS-CoV-2, was used as a control compound. In total, 10 μM of remdesivir significantly inhibited the replication and subsequent translation of the replicon, whereas dimethyl sulfoxide (DMSO) control did not (Figure 3A). The 50% inhibitory concentration (IC_50_) and 50% cytotoxicity concentration (CC_50_) values were calculated to 0.29 μM and more than 50 μM, respectively (selectivity index [SI] >172.4) (Figure 3B). The IC_50_ value estimated using our replicon system was about 2.6 times lower than the previously reported IC_50_ (0.77 μM) [11]. A previous study infected Vero E6 with infectious SARS-CoV-2 in the presence of remdesivir, and quantified the virus released in the supernatant by qRT-PCR at 48 hours post-infection [11]. The differences of our replicon assay and previous infectious SARS-CoV-2 assay including cell line (CHO or Vero), incubation time (24 h or 48 h), and action point of analysis (only RNA replication or whole replication steps) might cause the difference in IC_50_. Indeed, the difference of the cell line caused different IC_50_ values of remdesivir [12]. Nevertheless, the result was generally consistent with the previous report, thus demonstrating that our replicon system could be used for antiviral screening.

**Figure 3.**
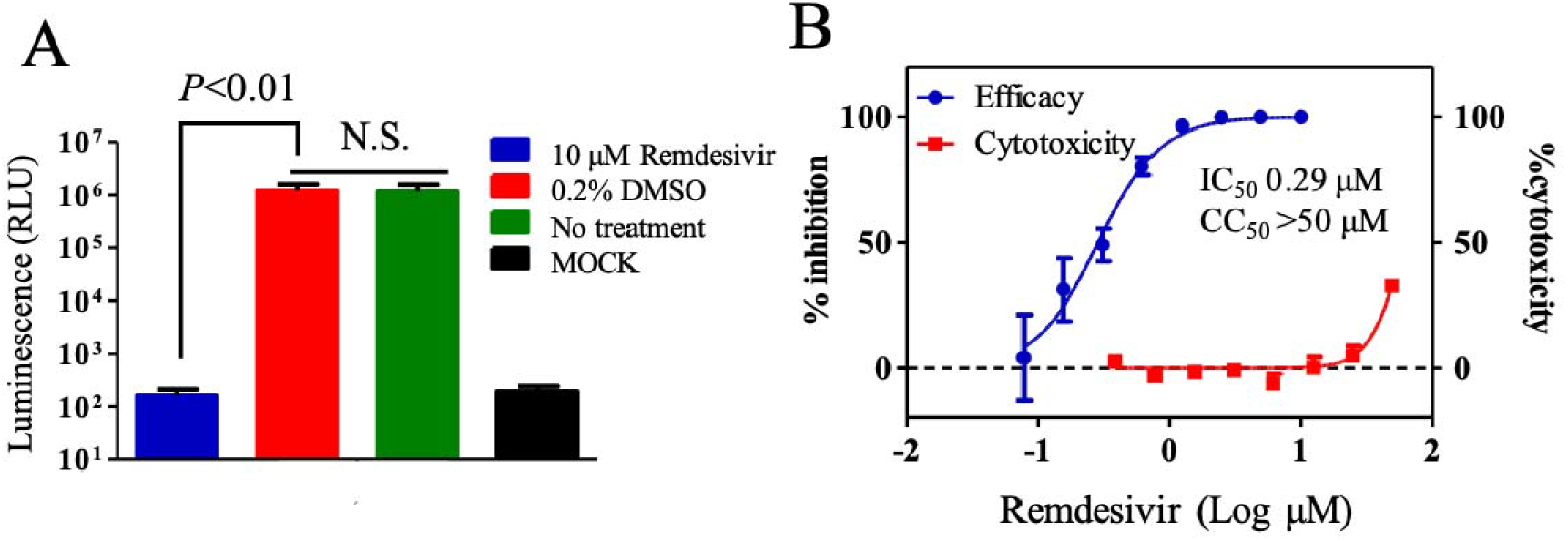
Antiviral evaluation using SARS-CoV-2 replicon. (A) Antiviral activity of remdesivir. The CHO-K1 cells electroporated with 5 μg of replicon RNA were seeded in a 96-well plate. The cells were treated immediately with 10-μM remdesivir or 0.2% DMSO. Luminescence was measured at 24 hours post-treatment. The mean and standard error of two independent experiments are shown in this figure. A one-way ANOVA was performed to determine the statistical significance. A *p*-value less than 0.05 was considerd to be statistically significant. N.S., not significant. (B) Calculation of IC_50_ and CC_50_. The CHO-K1 cells electroporated with replicon RNA was seeded. The cells were immediately treated with remdesivir at indicated concentrations. Luminescence and cell viability were measured at 24 hours post-treatment. IC_50_ and CC_50_ values were calculated by GraphPad software. The mean and standard error of two independent experiments are shown in this figure.

## Discussion

SARS-CoV-2 is an emergent threat worldwide. A high throughput and safe antiviral screening system is urgently needed to identify the anti–SARS-CoV-2 compound, which has not yet been developed. Here, we firstly reported a SARS-CoV-2 replicon system with PCR amplicon–based strategy. The advantage of this system is its technical simplicity. Additionally, this system enabled us to produce a replicon without generating genetically modified *E. coli*. Thus, this system can be handled even in a BSL-1 laboratory. Furthermore, bacteriotoxic elements in the SARS-CoV-2 genome do not affect the construction of the replicon. However, the PCR-based strategy might be inferior to the plasmid-based strategy in terms of the yield of replicon RNA and usability of genome modification. Additionally, PCR-based replicon might contain the undesired mutations, which are undetectable by Sanger sequence. Nevertheless, this PCR-based replicon system offered an alternative way over plasmid-based replicon, especially in the resource-limited settings.

In this study, BHK-21, 293T, and CHO-K1 cells were used because these cell lines were used for the construction of coronavirus replicon and coronavirus protein expression [4,5]. However, only CHO-K1 supported the robust replication of the replicon. The electroporation efficacy of large-size RNA to BHK-21 and 293T might be lower than that to CHO-K1 by the electroporation method used in this study. Alternatively, the host factors in BHK-21 and 293T cells might be related to the restriction of SARS-CoV-2 replication.

We chose to fuse HiBiT-tag to the N protein because subgenomic mRNA encoding N was the most abundantly produced mRNA during the replication of coronavirus [13]. This study demonstrated that the insertion of HiBiT-tag at the C-terminus of N protein did not disrupt the RNA replication. This finding could be applied to the construction of HiBiT-tagged reporter infectious virus [14]. We had also tried to fuse HiBiT-tag at the N-terminus of N protein. The luminescence of the replicon with N-terminal HiBiT was 10 times lower than that with C-terminal HiBiT at 24 hpt (supplementary Figure S1 and Table S1). The N protein is involved in not only nucleocapsid formation, but also RNA replication such as helicase activity and genome-length negative-strand RNA synthesis [15,16]. Although the N-terminus of N protein was not associated with either RNA binding or dimerization [17], the modification of the N-terminus might affect the replication efficacy.

This replicon system can be used not only for antiviral screening but also for the analysis of SARS-CoV-2 ORF1ab function in terms of RNA replication. SARS-CoV-1 replicon was applied to the functional analysis of non-structural proteins encoded in ORF1 [5]. Nowadays, several mutations have been observed in the replication complex regions because of worldwide pandemic [18]. For example, the virological meaning of ORF1ab 4715L mutation positively correlated to a high fatality rate remains unknown [19]. This system would help to shed light on the enigmatic SARS-CoV-2 RNA replication mechanism.

The disadvantages of this system were that our replicon was a transient expression system, which was not a high throughput system. The cell line stably carrying the replicon gene needs to be established by inserting the antibiotic resistance gene such as puromycin N-acetyl-transferase into the replicon genome [4]. Additionally, our replicon lacks the structural genes including S, E, and M. Thus, this system cannot be used for the compounds acting on receptor binding, virus entry, encapsidation, and virus release. These targets could be covered by using a single-round infectious pseudo-type reporter virus usable in the BSL-2 laboratory [20].

In conclusion, we reported a first SARS-CoV-2 replicon that can be applied to antiviral screening without using infectious virion. This replicon system would accelerate the antiviral screening and help to identify the novel drug candidates for COVID-19.

## Materials and Methods

### Virus and cell line

A clinical SARS-CoV-2 isolate from Japan (JPN AI-I 004 strain; EPI_ISL_407084) was used for the construction of replicon. Baby hamster kidney-21 (BHK-21) cell (ATCC: CCL-10) was maintained in the Eagle’s minimal essential medium (MEM) supplemented with 10% fetal bovine serum (FBS) at 37℃ with 5% CO_2_. Chinese hamster ovary-K1 (CHO-K1) cell (ATCC: CCL-61) was maintained in MEM supplemented with 10% FBS, non-essential amino acids at 37℃ with 5% CO_2_. HEK-293T cell (ATCC: CRL-3216) was maintained in the DMEM supplemented with 10% FBS.

### The construction of a SARS-CoV-2 replicon DNA

The viral RNA extracted from the culture fluid of SARS-CoV-2–infected Vero E6 cell (provided by the National Institute of Infectious Diseases, Japan) was reverse transcribed into cDNA by the SuperScript III First Strand Synthesis system (Thermo Fisher Scientific) with random hexamer primers. The fragments were amplified by primer sets (Table 1) and high-fidelity PCR with the Platinum SuperFi II DNA polymerase (Thermo Fisher Scientific). F8 was generated by the overlap PCR of F8A and F8B fragments to insert the HiBiT-tag at the C-terminus of N gene (Table 1). The overhang sequences after BsaI digestion were designed based on the ligase fidelity viewer program (available at the New England Biolabs website).

For assembly, all the fragments were digested with BsaI-HF v2 (New England Biolabs) and purified using NucleoSpin Gel and PCR clean-up (Macherey-Nagel). Then, two adjacent fragments of equimolar amount were mixed and ligated with 400 units of T4 DNA ligase (New England Biolabs) at 4℃ overnight: F1 (1.45 μg) and F2 (1.56 μg) for F1–2, F3 (0.86 μg) and F4 (0.85 μg) for F3–4, F5 (1.54 μg) and F6 (1.24 μg) for F5–6, and F7 (1.14 μg) and F8 (0.66 μg) for F7–8. The assembled fragments were electrophoresed on a 1% agarose gel and extracted using Monofas DNA extraction kit (GL Science). Then, extracted fragments were mixed and further assembled with 2,000 units of T4 DNA ligase at 4℃ overnight. The assembled DNA was directly purified by phenol–chloroform–isoamyl alcohol (25:24:1), by chloroform, and isopropanol precipitate. The pelleted DNA was washed once with 70% ethanol, dried by air, and finally dissolved in 10 μl of DEPC-treated water.

### RNA transcription, electroporation, and luminescence quantification

The replicon RNA was transcribed by the mMESSAGE mMACHINE T7 Transcription Kit (Thermo Fisher Scientific) according to the manufacturer’s instruction with some modifications. Cap analog to GTP ratio was set to 1:1. About 1 μg of the assembled DNA was subjected to RNA transcription. The reaction was incubated at 30℃ overnight. After removing the DNA template following the manufacturer’s protocol, RNA was extracted by phenol–chloroform and isopropanol precipitated. The pelleted RNA was washed once with 70% ethanol, dried by air, and dissolved in 40 μl of DEPC-treated water. The RNA was electrophoresed using DynaMarker RNA High for Easy Electrophoresis (BioDynamics Laboratory. Inc.) for the rough quality check.

The RNA was electroporated using NEPA21 electroporator (Nepagene). The cells were trypsinized and washed twice with Opti-MEM (Thermo Fisher Scientific). The washed cells (1 × 10^6^ cells) were mixed with 5 μg of replicon RNA in 100 μL of Opti-MEM. Electric pulses were given by NEPA21. The parameters for BHK-21 and CHO-K1 cells were as follows: voltage = 145 V; pulse length = 5 ms; pulse interval = 50 ms; number of pulses = 1; decay rate = 10%; polarity + as poring pulse and voltage = 20 V; pulse length = 50 ms; pulse interval = 50 ms; number of pulses = 5; decay rate = 40%; and polarity +/− as transfer pulse. The parameters for 293T cell was same as above except voltage 150 V and pulse length of 2.5 ms for poring pulse. After electroporation, the cells were seeded as 1.5 × 10^4^ cells/well in a 96-well plate. At various time points post-transfection, the cells were lysed with 25 μl of Nano-Glo HiBiT lytic detection system (Promega) plus 25 μl of PBS. The luminescence signal was detected by CentroPRO LB962 (Berthold Technologies).

### Immunofluorescence assay

At 24 hours post-transfection, the cells were fixed with 4% paraformaldehyde, followed by permeabilization with 0.5% Triton-X. After blocking with normal goat serum, the cells were incubated with primary mouse monoclonal antibodies (mAbs) (anti-N mAb [6H3: GeneTex] or anti-NSP8 mAb [5A10: GeneTex]) followed by a secondary antibody (goat anti-mouse IgG conjugated with Alexa Fluor 488). The cells were mounted in a mounting medium containing 4′,6-diamidino-2-phenylindole (DAPI: Vector Laboratories). Fluorescence images were acquired by a fluorescence microscope.

### Antiviral treatment

The CHO-K1 cells electroporated with 5 μg of the replicon RNA were seeded as 1.5 × 10^4^ cells/well in a 96-well plate. The cells were immediately treated with various concentrations of remdesivir. The cells were also treated with 0.2% DMSO as a negative control because 10-μM remdesivir contains 0.2% DMSO. At 24 hours post-treatment, the luminescence signal was detected as described above. Cell viability was measured by WST-1 assay following manufacture’s protocol (Roche). The IC_50_ and CC_50_ were calculated using a four-parameter logistic regression model from the GraphPad Prism 5 software (GraphPad Software Inc.).

## Acknowledgment

This work was supported by a young scientist dispatch program, Kobe University: by a subsidy for post-corona society realization, Hyogo prefecture, and by Japan Agency for Medical Research and Development (AMED) under Grant Number JP20he082206. The manuscript was proofread by Enago. We thank Ms. Honoka Yoneda for her technical assistance. We also thank Dr. Masayuki Saijo of the National Institute of Infectious Diseases for providing RNA of SARS-CoV-2 JPN AI/I 004 strain.

## Author contributions

T.K. and M.K. conceived the study. T.K. performed the experiments and took the lead in writing the manuscript. X.X., P. Y.-S., and M.K. provided feedback and helped shape the research and manuscript.

---

**Supplementary Table S1.**
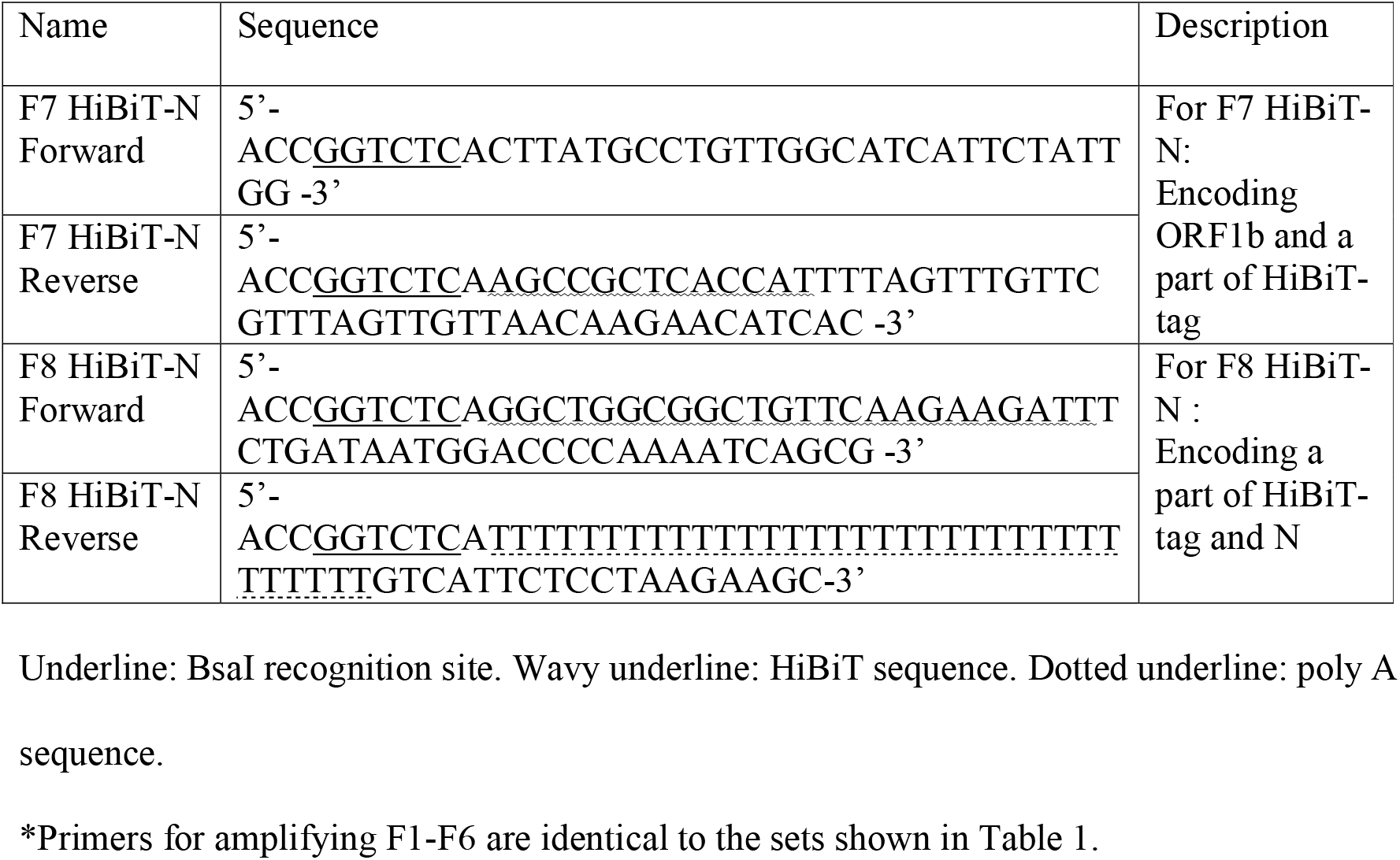
Primer sets for constructing a SARS-CoV-2 replicon with HiBiT-tag at the N-terminus of N protein*.

**Supplementary Figure S1.**
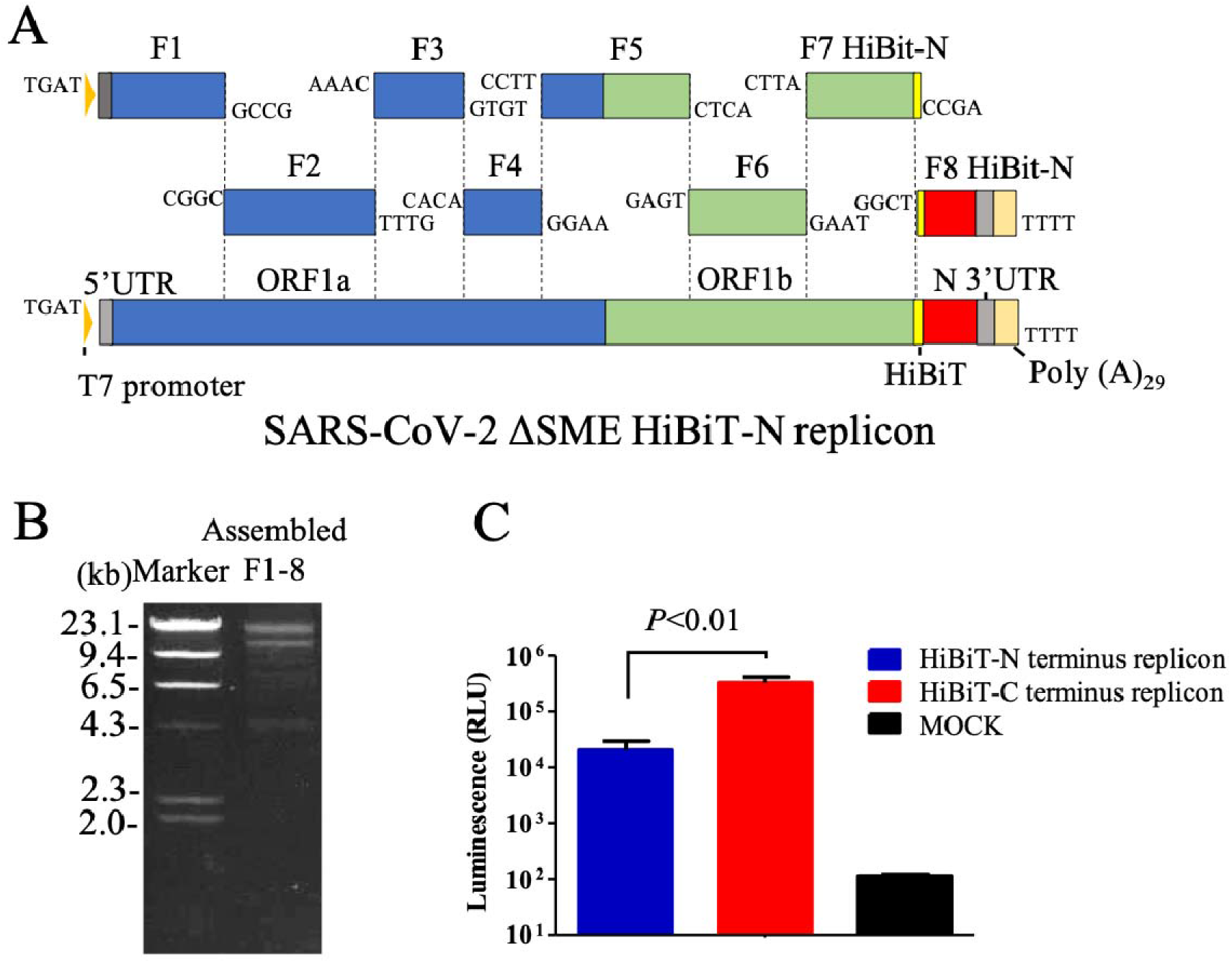
Construction and characterization of a SARS-CoV-2 replicon with HiBiT-tag at the N-terminus of N protein. (A) Strategy for *in vitro* assembly of a SARS-CoV-2 replicon DNA with HiBiT-tag at the N-terminus of N protein. The nucleotide sequences of the overhang are indicated in this figure. The replicon DNA was assembled using *in vitro* ligation. (B) Electrophoresis of an assembled DNA. About 100 ng of assembled DNA was run on a 1% agarose gel. The λ-HindIII digest marker is indicated in this figure. Successfully assembled replicon DNA was 23.2 kb. (C) Luminescence signals at 24 hpt. CHO-K1 cell was electroporated with 5 μg of replicon RNAs. Intracellular luminescence signals were measured at 24 hpt. The mean and standard error of two independent experiments are shown in this figure. A t test was performed to determine the statistical significance.

